# The rapid evolution of alternative splicing in plants

**DOI:** 10.1101/107938

**Authors:** Zhihao Ling, Thomas Brockmöller, Ian T. Baldwin, Shuqing Xu

## Abstract

Alternative pre-mRNA splicing (AS) is prevalent among all plants and is involved in many interactions with environmental stresses. However, the evolutionary patterns and underlying mechanisms of AS in plants remain unclear. By analyzing the transcriptomes of six plant species, we revealed that AS diverged rapidly among closely related species, largely due to the gains and losses of AS events among orthologous genes. Furthermore, AS that generates transcripts containing premature termination codons (PTC), although only representing a small fraction of the total AS, are more conserved than those that generate non-PTC containing transcripts, suggesting that AS coupled with nonsense-mediated decay (NMD) might play an important role in regulating mRNA levels post-transcriptionally. With a machine learning approach we analyzed the key determinants of AS to understand the mechanisms underlying its rapid divergence. Among the studied species, the presence/absence of alternative splicing site (SS) within the junction, the distance between the authentic SS and the nearest alternative SS, the size of exon-exon junctions were the major determinants for both alternative 5’ donor site and 3’acceptor site, suggesting a relatively conserved AS mechanism. Comparative analysis further demonstrated that variations of the identified AS determinants, mostly are located in introns, significantly contributed to the AS turnover among closely related species in both Solanaceae and Brassicaceae taxa. These new mechanistic insights into the evolution of AS in plants highlight the importance of post-transcriptional regulation in mediating plant-environment interactions.

**One sentence summary:** Changes of intron located splicing regulators contributed to the rapid evolution of alternative splicing in plants.

## INTRODUCTION

Due to their sessile lifestyle, plants have evolved various mechanisms to respond to environmental stresses. Alternative splicing (AS), a mechanism by which different mature RNAs are formed by removing different introns or using different splice sites (SS) from the same pre-mRNA, is known to be important for stress-induced responses in plants (Mastrangelo et al., 2012; Staiger and Brown, 2013). Both biotic and abiotic stresses such as herbivores (Ling et al., 2015), pathogens (Howard et al., 2013), cold (Leviatan et al., 2013) and salt (Ding et al., 2014) can all induce genome-wide changes in AS in plants. The environment-induced AS changes in turn can affect phenotypic traits of plants and may contribute to their adaptations to different stresses (Mastrangelo et al., 2012; Staiger and Brown, 2013). For example, low temperature-induced AS changes of flowering regulator genes affect flowering time and floral development in *A. thaliana* (Severing et al., 2012; Rosloski et al., 2013). The strong association between AS and environmental stimuli suggests that AS is involved in adaptation processes and thus evolved rapidly.

Two main functions of AS have been postulated: (i) to expand proteome diversity when different transcript isoforms are translated into different proteins (with different subcellular localizations, stability, enzyme activity etc.) (Kazan, 2003; Reddy, 2007; Barbazuk et al., 2008); (ii) to regulate gene expression by generating transcripts harboring premature termination codons (PTC) that are recognized by the nonsense-mediated decay (NMD) machinery and degraded (Chang et al., 2007; Kalyna et al., 2012; Kervestin and Jacobson, 2012). Although initially considered to be transcriptional noise, several AS events that introduce PTCs have been found to be highly conserved in animals (Ni et al., 2007; Lareau and Brenner, 2015) and plants (Iida and Go, 2006; Kalyna et al., 2006; Darracq and Adams, 2013), suggesting that that the combination of AS with NMD might play an important role in controlling mRNA levels post-transcriptionally. However, it is unclear whether NMD-coupled AS is more conserved than the AS that generates transcripts without PTC at a genome-wide level.

The evolution of AS in plants, compared to that in vertebrates, remains largely unclear. Studies that compared organ-specific transcriptomes from different vertebrate species spanning ~350 million years of evolution showed that AS complexity differs dramatically among vertebrate lineages, and AS evolved much faster than gene expression has (Barbosa-Morais et al., 2012; Merkin et al., 2012). For example, within 6 million years, the splicing profiles of an organ are more similar to other organs of the same species than the same organ in other species, while the expression profiles of the same organ are similar to the organ in other species (Barbosa-Morais et al., 2012; Merkin et al., 2012). In plants, largely due to the lack of comprehensive transcriptomic data, such comparative analysis remains unavaliable. However, several indications suggest that AS in plants and vertebrates may share evolutionary pattern of the rapid divergence. For example, only 16.4% of AS between maize and rice, and 5.4% between *Brassica* and *Arabidopsis* are conserved (Severing et al., 2009; Darracq and Adams, 2013). A more recent study further showed that only 2.8% of genes showed conserve AS between two species of mung beans, *Vigna radiate* and *V. angularis* (Satyawan et al., 2016). Furthermore, large changes in AS also exist between different ecotypes of the same species (Streitner et al., 2012). However, such low conservations of AS found among species could also be due to several other confounding effects. For example, it is also known that the levels of gene expression, which are highly associated with AS, also diverge rapidly in plants (Yang and Wang, 2013). As a consequence, it remains unclear whether the low observed levels of AS conservation results simply from the rapid expression changes between species. Furthermore, AS detection of is highly dependent on sequencing depth and the tissue types used for generating transcriptomic data (Xu et al., 2002; Ellis et al., 2012; Ling et al., 2015). Therefore, it is necessary to systematically control for different confounding effects in order understand the evolutionary pattern of AS in plants.

From a mechanistic perspective, divergence of AS among species is contributed by factors that affect the exon-intron splicing process, which is mediated by the spliceosome. While the recognition processes of exonic and intronic regions are directed by sequence features of the pre-mRNA in animals, how the spliceosome removes introns and ligates exons is poorly understood in plants. In metazoans, it is known that four crucial signals are required for accurate splicing: (i) 5’ splice sites (SS), which contains a GU dinucleotide at the intron start surrounded by a piece of longer consensus sequences of lower conservation, (ii) 3’ SS, which includes an AG at the 3’ end surrounded by similar sequences of 5’ SS, (iii) a polypyrimidine tract (PPT) and (iv) a branch site (BS) sequence located ~17-40 nt upstream of the 3’ SS (Black, 2003; Fu and Ares, 2014). In plants, similar sequence features with a small difference at specific positions were found, except for the requirement of a BS (Reddy, 2007). In addition, a UA-rich tract in introns has also been found to be important for efficient splicing in plants (Lewandowska et al., 2004; Simpson et al., 2004; Baek et al., 2008). In animals, the regulation of splicing also depends largely on *cis* signals and *trans*-acting splicing factors (SFs) that can recognize the signals (Barbosa-Morais et al., 2012; Merkin et al., 2012). Among different SFs, serine/arginine-rich (SR) proteins are from an important SFs family that has been shown to be involved in AS regulation (Lopato et al., 1999; Gao et al., 2004; Wang and Brendel, 2004; Reddy, 2007; Reddy and Shad Ali, 2011). In addition, many splicing regulatory elements (SREs) and RNA-binding proteins (RBPs) have been identified in animals, and the interactions among these SREs in the pre-mRNA and RBPs were found either to promote or suppress the use of a particular splice sites (Licatalosi et al., 2008; Chen and Manley, 2009; Barash et al., 2010). The number of SR proteins genes in plants is nearly twice that of number found in non-photosynthetic organisms, although the number varies among different species (Iida and Go, 2006; Isshiki et al., 2006; Richardson et al., 2011). To date, more than 200 RBPs and 80 SREs in plants have been identified using computational approaches (Lorkovic, 2009), however, only a few of these have been functionally validated (Yoshimura et al., 2002; Pertea et al., 2007; Schoning et al., 2008; Thomas et al., 2012).

In mammals, the emergence of AS originated from constitutive splicing with the fixation of SREs and the creation of alternative competing SS (Koren et al., 2007; Lev-Maor et al., 2007). Distinctive features that distinguish alternatively spliced exons/introns from constitutively spliced exons/introns can be used to accurately predict the specific AS type (Koren et al., 2007; Braunschweig et al., 2014). Furthermore, other factors including secondary and tertiary RNA structures, chromatin remodeling, insertion of transposable elements (TEs) and gene duplication (GD) may also be involved in regulating AS (Liu et al., 1995; Sorek et al., 2002; Donahue et al., 2006; Su et al., 2006; Kolasinska-Zwierz et al., 2009; Schwartz et al., 2009; Warf and Berglund, 2010; Lambert et al., 2015). However, the extent to which changes in these factors contributed to the evolutionary history of AS in vertebrates remains largely unclear. Recently, a study using millions of synthetic mini-genes with degenerated subsequences demonstrated that the likelihood of AS decreases exponentially with increasing distance between constitutive and newly introduced alternative SS (Rosenberg et al., 2015), suggesting that sequence changes between constitutive and alternative SS might contribute to the rapid changes of AS among species. In plants, however, the detailed mechanisms that affect AS remain largely unclear (Reddy et al., 2013). Although it has been proposed that changes in chromatin features such as DNA methylation, histone marks as well as RNA structural features, and SREs are important in regulating AS in plants, experimental evidence remains largely lacking (Reddy et al., 2013). A recent study shows that DNA methylation could affect AS in rice (Wang et al., 2016), indicating changes in DNA methylation can contribute to the variations of AS among species, however, this hypothesis has not been thoroughly tested.

Because AS regulation is a complex process involving many factors, computational modeling is a useful tool for identifying key factors and predicting the outcome of splicing. While the Bayesian neural network (BNN) method was developed for decoding the splicing code in mammals (Barbosa-Morais et al., 2012), deep learning approaches, which refers to methods that map data through multiple levels of abstraction, have recently been shown to surpass BNN-based approaches(Leung et al., 2014; Mamoshina et al., 2016). Furthermore, deep learning methods are also able to cope with large, heterogeneous and high-dimensional datasets and problems that include predicting DNA and RNA-binding proteins (Alipanahi et al., 2015) and AS (Leung et al., 2014; Mamoshina et al., 2016).

Here we performed a comparative analysis of the transcriptomes of both closely and distantly related plant species to explore the evolutionary history of AS in plants. To further understand the mechanisms underlying the AS evolution in plants, we applied a deep learning approach to investigate the determinants of AS and their effects on AS evolution. Specifically, we aimed to address the following questions in plants: 1) What are the evolutionary pattern of AS? 2) Are the AS events that are coupled with NMD more conserved than regular AS events? 3) Which factors are important in determining AS? 4) Which factors have contributed to the rapid turnover of AS between closely related plant species?

## RESULTS

### Genome-wide AS patterns are species-specific in plants

To provide an overview of AS evolution among different plant families, we studied the genome-wide AS in *Arabidopsis thaliana*, soybean (*Glycine max*), tomato (*Solanum lycopersicum*) and wild tobacco (*Nicotiana attenuata*), for which comparable transcriptomic data sets are available from the same tissues (roots, leaves and flowers) and represent a wide-range of eudicots. The overall distributions of different AS types within each species are consistent with previous studies. In all investigated species, intron retention (IR) and alternative 3’ acceptor site (AltA) are the two major AS types (Figure S1) (Aoki et al., 2010; Marquez et al., 2012; Shen et al., 2014; Ling et al., 2015).

To investigate the evolutionary patterns of AS, we compared AS profiles across selected tissues and species. Because sequencing depth is known to strongly affect AS detection, we randomly subsampled 16 million (the lowest depth among all samples) uniquely mapped reads from each sample to standardize for the heterogeneity of sequencing depths. All downstream comparative analyses were based on this subsampled dataset. Clustering analyses using percent spliced index (PSI) that measures the qualitative differences of AS among samples showed that different tissues of the same species are more similar to each other than the same tissue from different species (Figure 1A). This pattern is consistent with the hypothesis that AS evolves rapidly in plants. Using qualitative measures of AS that consider the presence or absence of AS (binary) for all one-to-one orthologous genes, the same species-specific clustering pattern was found (Figure 1B). Furthermore, consistent results were also obtained when each type of AS was analyzed separately (Figure S2).

**Figure 1.**
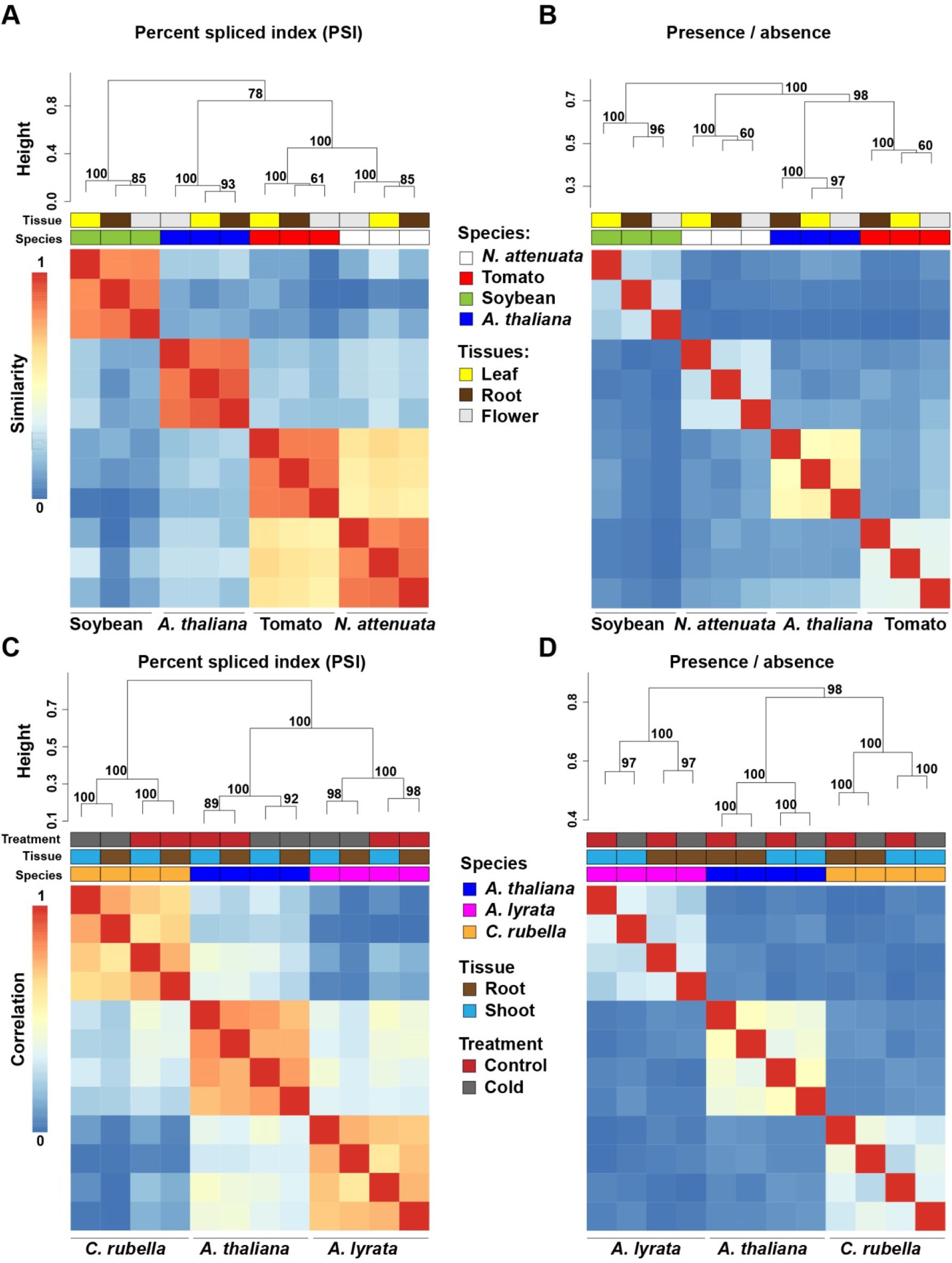
Species-specific clustering of alternative splicing (AS) among different plant species. **(A)** and **(C)**, heatmaps depict species-specific clustering based on PSI values amongfour eudicots species **(A)** and three Brassicaceae species **(C)**. The clustering is based on conserved splicing junctions (A and C: n = 502 and 5241, respectively). **(B)** and **(D)**, heatmaps depict species-specific clustering based on presence and absence of AS of the one-to-one orthologous genes. In total, junctions from 3,857 **(B)** and 6,262 **(D)** orthologous were used for the clustering. Numbers present in each branch node represent the approximately unbiased bootstrap value calculated from 1000 bootstrap replications. The color code above each heatmap represents species, tissue, and treatments.

To further investigate the evolutionary patterns of AS among closely related species, we analyzed a recently published transcriptome dataset from three Brassicaceae species (*A. thaliana*, *Arabidopsis lyrata* and *Capsella rubella*), each of which have comparable transcriptome data from two tissues (root and shoot) and two treatments (control and cold treated) (Seymour et al., 2014). Using both quantitative (PSI) and qualitative measures (binary) of AS, a similar species-specific clustering pattern was observed (Figure 1C and D). Interestingly, within same species and same tissue, samples exposed to cold stress clustered together regarding levels of PSI, a result which is consistent with previous studies that demonstrate that stresses can induce genome-wide AS responses (Li et al., 2013; Ding et al., 2014; Ling et al., 2015).

### Genome-wide AS regulations diverge faster than gene expression among closely related species

Species-specific clustering patterns were also reported at the level of gene expression (GE) among *A. thaliana*, rice and maize (Yang and Wang, 2013). To examine whether species-specific AS clustering results from gene expression divergences, we compared the divergence patterns of AS and GE among transcriptomes of different species. Comparisons among species from different plant families showed that both GE and AS cluster in species-specific patterns (Figure 1A and B, S3A and B). However, when species from the same plant family are compared, such as tomato and *N. attenuata* (Solanaceae), the species-specific AS pattern remained (Figure 1A and B), but the GE data clustered in tissues-specific pattern (Figure S3A and B). This suggests that the expression profiles of the same tissues from different species are more similar to each other than the expression patterns from different tissues of the same species. This result was also found in the expression profiles of tissue samples from the three Brassicaceae species, among which the expression profiles of shoots and roots from different species were clearly separated (Figure S3C and D). These results indicate that AS evolves faster than GE in plants, the pattern of which is similar to that in animals (Barbosa-Morais et al., 2012; Merkin et al., 2012).

### Rapid gains and losses of AS among different species

Species-specific clustering of AS pattern suggests low level of AS conservation among species. Overall, among 3,857 one-to-one orthologous genes among the four dicots that have AS in at least one species, only ~7% of them have AS in all four species, while ~41% of them have species-specific AS. The rapid change of AS could result from the rapid loss or gain of EEJ between species. To exam the conservation of EEJs, we compared the EEJ structures among pairwise orthologous genes. In total, 60% of EEJs are conserved in at least two species (while only ~12% for AS), and the analysis based on AS events from the most conserved EEJs (found in all four species) showed that 92% of these are species-specific (Figure S4A). A similar pattern was found based on the analysis of EEJs that are conserved in at least two species, in which only 10 AS events (0.25%, out of 4,015) were found conserved among all four species (Figure S4B). We also performed the same analysis within the three Brassicaceae species and found 69% of total EEJs are conserved in at least two species (while only 27% of AS are conserved) and 72% of AS events that were found at the EEJs shared among all three species were species-specific (Figure S5A). Furthermore, only ~8% of AS events (1,476 out of 19,170) are conserved among all three species (Figure S5B). Together, the results between divergent species and closely related species consistently suggest that AS diverge rapidly in plants.

To investigate the transition spectrum of AS at the conserved EEJs between each species pairs, we calculated the transitions among different types of AS. Among the four dicots, while the transitions among different AS types are rare, the gain/loss of AS was found to be the most abundant transition type among all comparisons (Figure 2A-F). Among different AS types, AltA and ES are the most and least conserved AS, respectively. Similar patterns were observed among three closely related species in Brassicaceae (Figure S6A-C). These results suggest that the species-specific AS pattern is largely not due to the rapid changes of EEJs among species, but rather the rapid species-specific gains and losses of AS.

**Figure 2.**
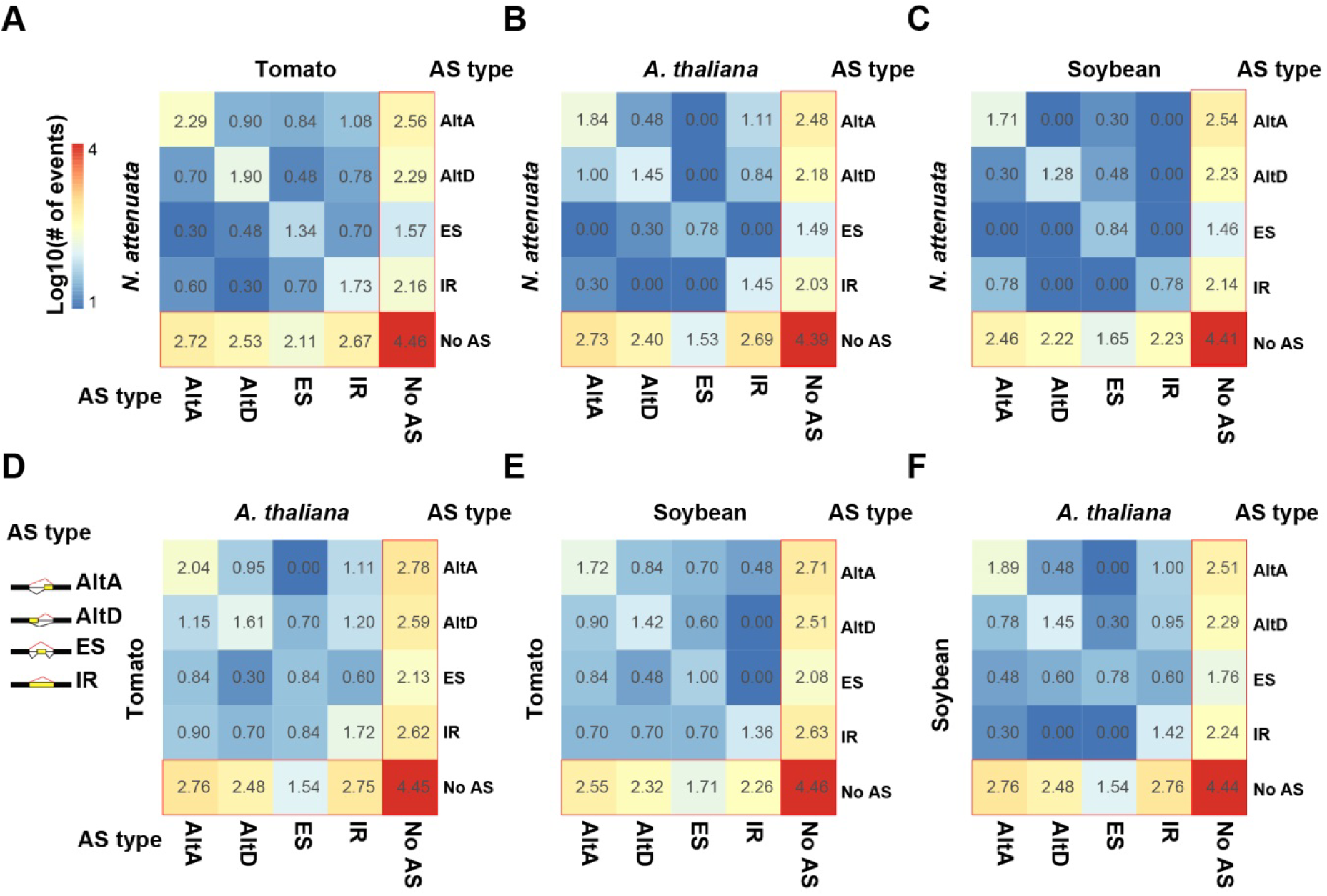
Transition spectrum of AS between each species pairs. **(A)** *N. attenuata* vs tomato, **(B)** *N. attenuata* vs *A. thaliana*, **(C)** *N. attenuata* vs soybean, **(D)** tomato vs *A. thaliana*, **(E)** tomato vs soybean and **(F)** soybean vs *A. thaliana*. The color of each grid refers to log10 transformed number of AS events. The transformed values are also shown in the middle of each grid. AltA: alternative 3’ acceptor site; AltD: alternative 5’ donor site; ES: exon skipping; IR: intron retention.

### The group of AS that generate PTC-containing alternative transcripts is more conserved than others

Previous studies suggest that many pre-mRNAs underwent unproductive AS, which generates transcripts with in-frame PTCs that are coupled with nonsense-mediated decay (NMD) in plants (Kalyna et al., 2012; Drechsel et al., 2013). To investigate whether unproductive AS can affect the AS conservation and contribute to the rapid loss/gain of AS among different plant species, we separated the AS into two groups: (1) PTC+ AS and (2) PTC- AS (details see Materials and Methods). Overall, the portion of PTC+ AS ranges from 9% - 15% among the four dicots (Figure S7), suggesting that only a small portion of AS generated PTC-containing transcripts. Comparing the levels of conservation between tomato and *N. attenuata*, we found the PTC+ AS is significantly more conserved than PTC- AS (*P* < 0.02, Figure 3A). For example, among nine PTC+ AS of *N. attenuata* which are both conserved and have PTC information in tomato, eight of them (89%) also generated PTC+ transcripts in tomato.

**Figure 3.**
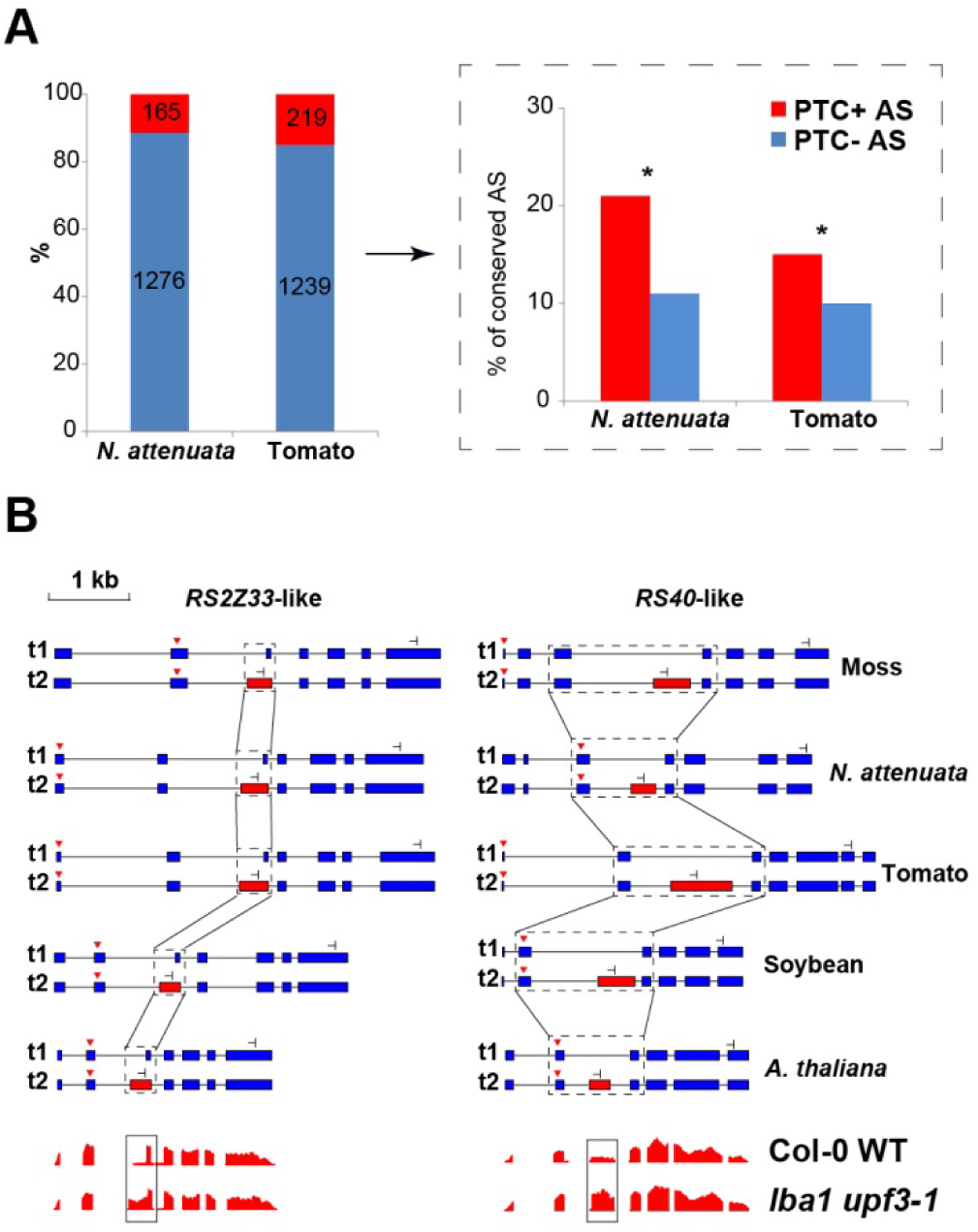
The conservation of AS between PTC+ and PTC- AS. **(A)** The number and relative portions of PTC+/- AS in *N. attenuata* and tomato. The insert indicated by the black arrow depicts the likelihood of PTC+ and PTC- AS that are conserved between *N. attenuata* and tomato. Asterisks indicate the significance as determined by Fisher’s exact test (P < 0.05). **(B)** Conserved AS between moss and eudicots in serine/arginine-rich splicing factor *RS2Z33*-like and *RS40*-like gene. The diagrams of the structure of transcripts generated by the AS in all five species, the dominant and minor transcripts are represented by t1 and t2, respectively. Constitutive exons are represented by blue boxes, alternatively spliced exons are represented by red boxes and introns are represented by black solid lines. The black dotted boxes highlight the conserved AS and the start and stop codons are shown as red triangles and stop signs, respectively. The diagrams in the bottom panel showed the relative read coverage of *AtRS2Z33* and *AtRS40* exons in wild type plant and *lba1 upf3-1* double mutants. The black box highlights the coverage of the spliced region which is significantly increased in *lba1 upf3-1* double mutants (The diagrams are modified based on the data shown in http://gbrowse.cbio.mskcc.org/gb/gbrowse/NMD201)

To further investigate the level of conservations of PTC+ AS, we extended our analysis by adding the transcriptome data of a very ancient plant species, the spreading earth moss (*Physcomitrella patens*). We focused on the 10 most highly conserved AS events found in all four dicot plants (Figure S4B) and checked their presence in moss. In total, we found six AS events that were also present in moss, indicating these AS events might have evolved since land plants and played essential functions in plants. Interestingly, two of these ultra-conserved AS events were from serine/arginine-rich (SR) genes (*RS2Z33*-like and *RS40*-like), which are part of RNA splicing machinery and the *RS2Z33*-like gene also has AS in rice (Iida and Go, 2006). Analyzing the protein coding potential of the transcripts generated by these six ultra-conserved AS events showed that five resulted in PTC+ transcripts. For example, the AS events of *RS2Z33*-like and *RS40*-like genes result in PTC+ alternative transcripts in all five species and are likely the targets of NMD (Figure 3B). To further investigate whether these PTC+ transcripts are regulated by NMD, we analyzed the available transcriptome data from *A. thaliana* wild type (WT) and NMD-deficient (*lba1* and *upf3-1* double mutant) plants (Drechsel et al., 2013). Among all five PTC+ transcripts in *A. thaliana*, three showed significantly higher expression in NMD-deficient plants (Drechsel et al., 2013) (*P* < 7e-06), including *RS2Z33*-like and *RS40*-like genes (Figure 3B). Together, these results suggest that AS coupled with PTC is more conserved than regular AS and some of these AS-PTC pairs may play essential roles in plants.

### Mechanisms involved in determining AS are overall conserved among different plant species

To further understand the mechanisms that contributed to the rapid turnover of AS among species, it is necessary to identify the key features of AS in plants, which was largely unknown (Reddy et al., 2013). Because splicing is often mediated by SS, we were interested in whether the SS were different between constitutively and alternatively spliced junctions. Comparisons of the SS and their surrounding 12 bp sequences between constitutively and alternatively spliced junctions revealed that their SS are overall very similar (Figure S8). Furthermore, we separately identified sequence motifs (12-mer) that are enriched in 5’ and 3’ splice sites (SS) compared to random sequences and found that these identified motifs are also highly conserved among studied species (Figure S9).

When alternative SS is present, the distance between the regular and the nearest alternative SS and inter-GT/AG, splicing junction size and the strength of the alternative SS are also important for the regulation of different AS types (Gopal et al., 2005; Kandul and Noor, 2009; Braunschweig et al., 2014; Rosenberg et al., 2015). For the different AS types, we compared these features from both constitutively and alternatively spliced junctions. Because exon skipping (ES) events are rare in all species, we only studied the three most abundant AS types (AltD, AltA and IR). The results showed that for a given junction, while the likelihood of both AltD and AltA decreases with the distance between authentic and alternative SS as well as the distance between authentic SS and the nearest internal GT/AG, the likelihood of both AltD and AltA increases with junction size (Figure 4A and B). Interestingly, although the likelihood of IR in smaller junctions appears larger than in large junctions, no significant correlation with junction size was found (Figure S10A). Both 5’ and 3’ SS of the junction with IR are significantly weaker than constitutive junction (Figure S10B).

Furthermore, the presence/absence of UA-rich tract, polypyrimidine (PPT) tract, branch site (BS) is also known to be associated with 3’ splicing recognition in eukaryotes (Lewandowska et al., 2004; Fu and Ares, 2014). We compared the frequency of AltA and IR between junctions of the AS gene with and without the presence of UA, PPT tract and BS within 100 bp upstream of 3’ SS. We found the frequencies of both AltA and IR are significantly higher in junctions without UA and PPT than junctions with them, while the presence of BS had no significant effect (Figure S11).

**Figure 4.**
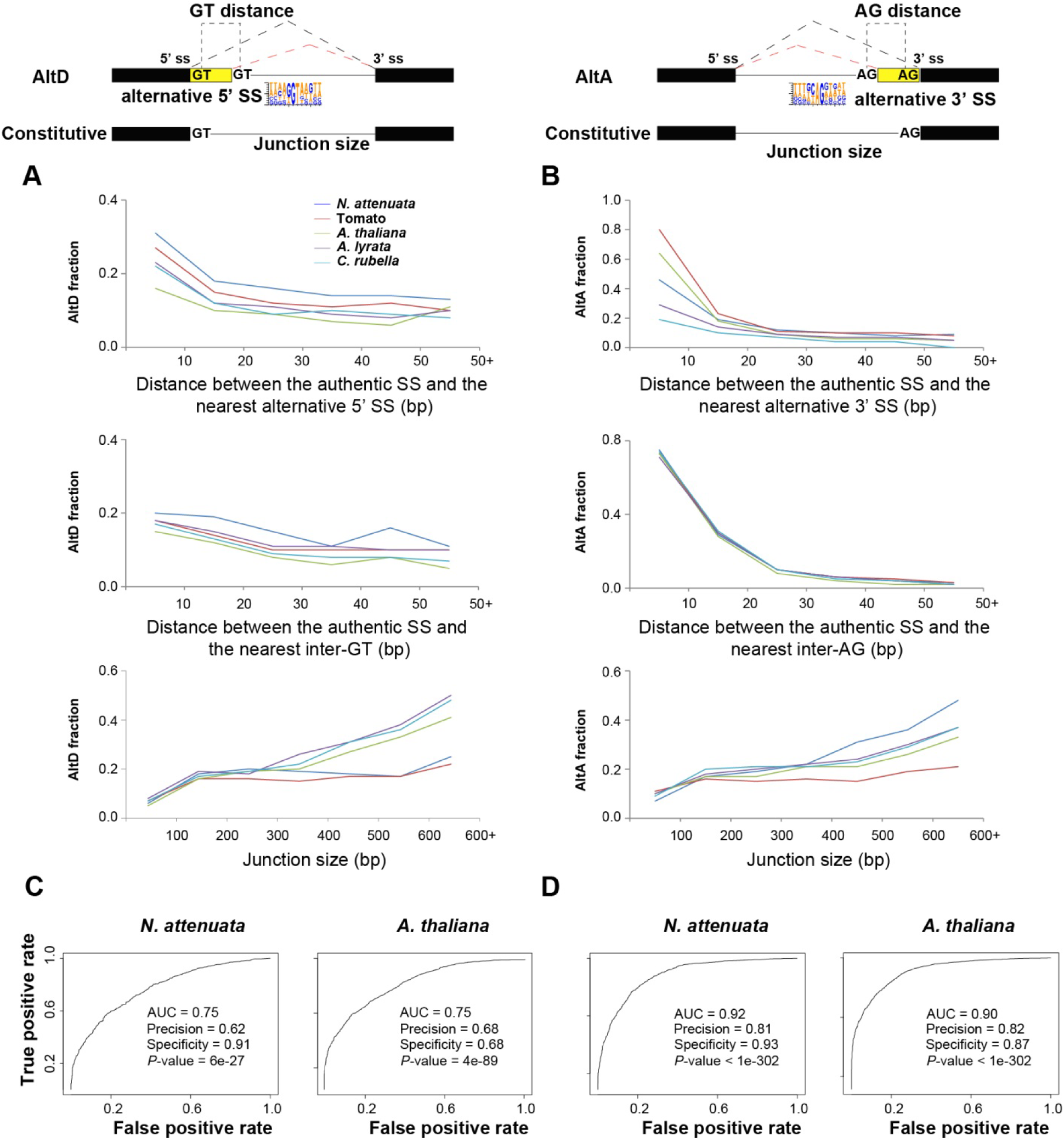
The determinants of alternative 5’ donor site (AltD) and alternative 3’ acceptor site (AltA) in plants. **(A)** and **(B)**, the frequencies of AltD/AltA on junctions with different distance between the authentic SS and nearest alternative SS (5’ ss and 3’ ss, respectively), and the distance between authentic SS and the nearest inter GT/AG and junction size. **(C)** and **(D)**, the area under the curve (AUC) plot of deep learning models using the key determinants of AltDand AltA in *N. attenuata* and *A. thaliana*. The model performance including area under the curve (AUC), accuracy, specificity and significance are also shown.

*Cis*-regulatory elements, including enhancers and silencers near SS are also important for the regulations of splicing. To identify these candidate regulatory elements, we performed *de novo* hexamer motif enrichment analysis by comparing 50 bp sequences from 5’ and 3’ sides of both donor and acceptor sites between alternatively spliced and constitutively spliced junctions. The results showed that most of the putative enhancer motifs for alternatively spliced junctions are highly similar to the identified SS. In addition, we also identified several putative silencer motifs (range from 5-10 for AltD and 10-18 for AltA in the five species), which were significantly more enriched in constitutively spliced junctions than alternatively spliced junctions (Figure S12 A and B).

To evaluate whether these identified features represent the AS determinants, we used a machine learning approach and modeled the different types of AS in each of the studied species. The rationale for this approach is that if the features we identified as representative of the key AS determinants were accurate, we would be able to predict whether an exon-intron junction is constitutively or alternatively spliced based on their quantitative or qualitative information. For this, we combined all of extracted featured mentioned above. In addition, we also extracted information on whether the alternative SS would introduce a frameshift, which may result in premature terminate code (PTC) or open reading frames, the number of reads that support the junction, which represent levels of expression that is known to be associated with AS, as well as the presence and absence of the identified *cis*-motifs. Using this information, our model achieved high precision and specificity for both AltD and AltA in all five species (Figure 4C and D, S13A and B), which suggests that the identified features can provide sufficient information to discriminate AS junctions from constitutively spliced junctions. However, for IR, the extracted features were not predictively useful (the average AUC is 0.54), indicating that additional undetected factors have contributed to the determination of IR.

This modeling approach further provides indicative information on the relative importance of each feature to the prediction model. The results showed that for AltD, the distance between the authentic SS and the nearest alternative 5’SS or inter GT, the junction size and presence/absence of 5’ additional SS in the intron are among the top important features for the prediction in all species (Supplemental file 1). In addition, the frame shifts introduced by the nearest alternative 5’ SS and nearest GT were also important contributors to the model (Supplemental file 2). For AltA, the distance to the nearest inter-AG dinucleotide is the top feature for the prediction among all five species. Interestingly, all of the identified putative silencers only had a marginal role for the predictions of both AltD and AltA (Supplemental file 2). Together, these results showed that the mechanisms regulating AltD and AltA are likely overall conserved among the studied species.

### Changes in AS determinants contributed to the rapid turnover of AS in plants

The relatively conserved AS regulation mechanisms among studied species provide a foundation for investigating the mechanisms that contributed to a rapid turnover of AS among closely related plant species. We hypothesized that the changes in the identified AS determinants among species resulted in a rapid divergence of AS in plants. To test this, we associated the changes of the identified AS determinants and AS conservation among closely related species. Because we did not find determinants for IR, we only focused on the evolution of AltA and AltD.

Variation in the distance between authentic SS and alternative SS or inter-GT/AG were negatively associated with AS conservation: the levels of AS conservation decreased with increasing distance in all three pairs of comparisons (Figure 5A and B), for both AltD and AltA. In addition, the changes in the reading frame introduced by the alternative SS also significantly decreased the conservations of both AltA and AltD (Figure 5C and D). The similar pattern was also found for the distance between authentic SS and the nearest inter-GT/AG (Figure 5E-H).

**Figure 5.**
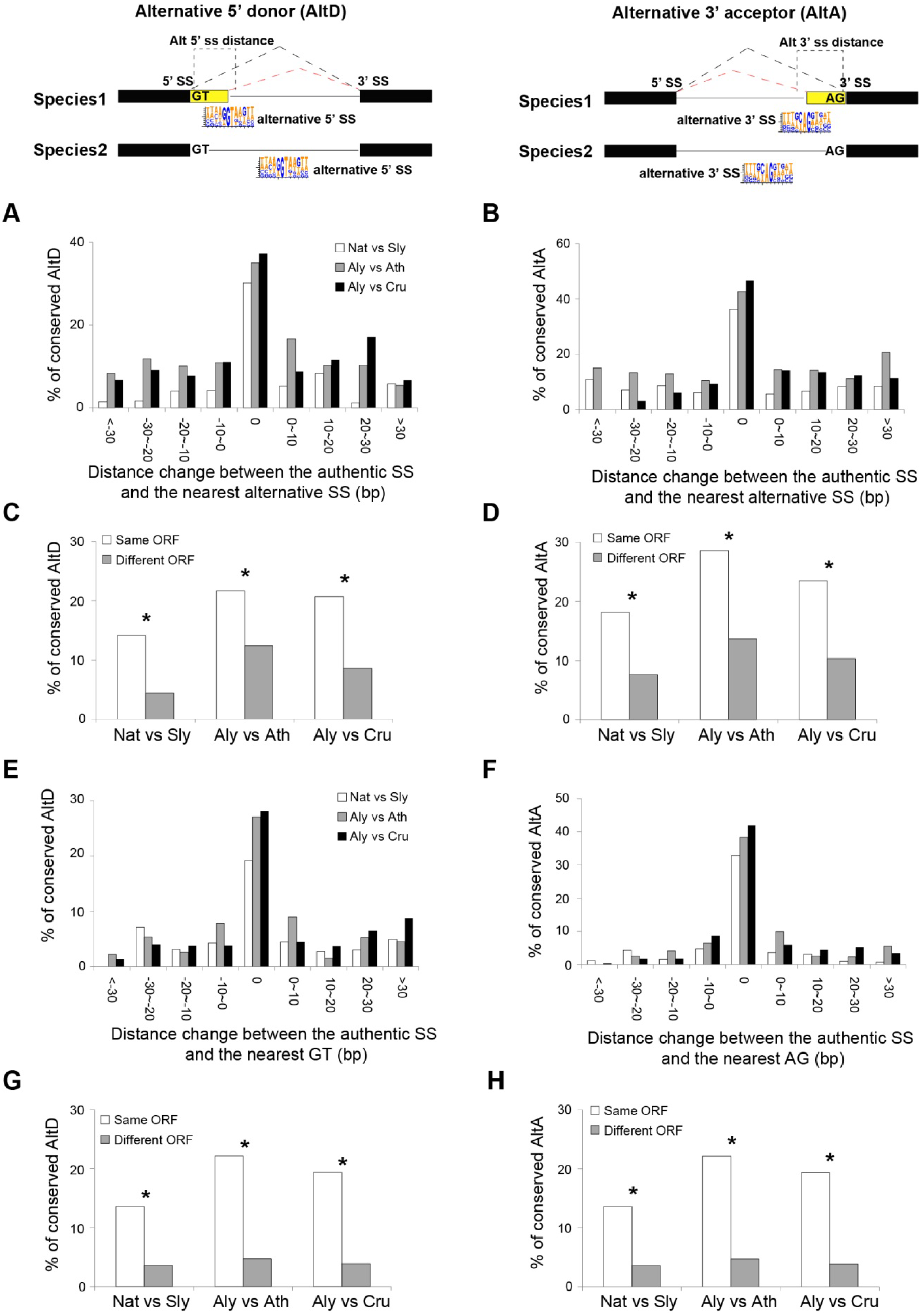
Features that affect the conservation of AltD and AltA between closely related plant species. **(A)** and **(B)**, the portion of conserved AltD/AltA decreases with changes of distance between authentic and alternative SS between two species. **(C)** and **(D)**, the percent of conserved AltD/AltA in the group that the nearest alternative 5’/3’ SS in the two species generate the same or different ORF transcripts. **(E)** and **(F)**, the portion of conserved AltD/AltA decreases with changes in the distance between authentic SS and nearest inter-GT/AG sites between two species. **(G)** and **(H)** the percent of conserved AltD/AltA in the group that the nearest inter-GT/AG in the two species generate transcripts with same or different open reading frame (ORF). Nat: *N. attenuata*, Sly: Tomato, Ath: *A.thaliana*, Aly: *A. lyrata*. The asterisks indicate the significance as determined by Fisher's exact test (*P* < 0.05).

Variation in the *cis-*regulatory elements (CREs) UA-tract, PPT and BS (Figure S11 and Supplemental file 2) significantly reduced the conservation for AltA (Figure S14B), but did not affect the conservation of AltD among species (Figure S14A). This result is consistent with the functional roles of these CREs in regulating AltA.

To further systematically analyze different factors that might affect the conservation of AS, we constructed an AS evolution model for each closely related species pair using a deep learning method. In addition to the key AS determinants identified in this study, we also included several other features that were previously hypothesized to be important for AS conservation between species in the model, such as changes in copy numbers (role of gene duplications), transposable element (TE) insertion within the junction, GC-content and sequence similarity of SS. For the AltD, all three models between species pairs achieved significantly better prediction than by chance (highest *P*-value = 3e-44), with an average precision of 0.63 and specificity of 0.82. In all three pairwise comparison models, the distance changes between authentic and nearest alternative 5’ SS or inter-GT/AG are among the top five important features (Figure S15A and Supplemental file 1). For AltA, all three models achieved a precision and specificity (average 0.70 and 0.85, respectively) that was significantly higher than by chance (highest *P*-value = 3e-145). In all three models, distance changes between authentic SS and the nearest inter-AG or alternative 3’ SS and the changes on CREs (UA and PPT tracts) represent the top five most important features that contributed to the model predictions (Figure S15B and Supplemental file 1).

Interestingly, we found TE insertions to also be an important factor that reduced the conservation of both AltD and AltA between *N. attenuata* and tomato but not between any of two Brassicaceae species (Figure S15A and B). This is likely due to the difference of TE abundance between *N. attenuata* (~63%) and tomato (~81%), values which are much higher than the differences between *A. thaliana* (~23%) and *A. lyrata* (28%) (Hu et al., 2011; Tomato Genome, 2012). Furthermore, we also analyzed the impact of DNA methylation changes between *A. thaliana* and *A. lyrata* using data from (Seymour et al., 2014) and found no significant effects (Figure S15).

## DISCUSSION

Here we showed that alternative splicing (AS) diverges more rapidly than does gene expression (GE) and the rapid gain and loss of AS resulted in lineage-specific AS profiles in plants. Although AS events that introduce premature termination codons (PTC), represent only a small proportion of the total AS events, they are more conserved than AS events that do not introduce PTC (Figure 3A). Consistently, several AS events that generate PTC-containing transcripts were ultra-conserved among highly divergent plants. To understand the mechanisms that resulted in a rapid turnover of AS between closely related species, we identified several key determinants for both alternative donor (AltD) and alternative acceptor (AltA) splicing, and found that the change of these key determinants between species significantly contributed to the rapid gain and loss of AS in plants.

In this analysis, we observed a dominant species-specific pattern of AS among different species, suggesting that AS in plants diverges rapidly (Figure 1). Such rapid evolution of AS in plants is similar to the pattern found among vertebrate species that span ~350 million years of evolution (Barbosa-Morais et al., 2012; Merkin et al., 2012), in which AS is largely segregated by species, while GE is segregated by tissue types (Barbosa-Morais et al., 2012; Merkin et al., 2012). This indicates that the rapid evolution of AS might be universal among eukaryotes. Interesting, in plants, the evolution of GE appears to be faster than in vertebrates, as the tissue-dominant clustering of GE was only observed among closely related species, but not among species from different families (Figure S3). This pattern is consistent with a previous study which showed the overall GE of three highly divergent species, including both monocots and dicots (diverge ~200 million years ago), are grouped according to species rather than organs (Yang and Wang, 2013). In vertebrates, some tissues, such as brain, testis, heart and muscle still showed a strong tissue-specific splicing signature, despite the dominant species-specific splicing background (Barbosa-Morais et al., 2012; Merkin et al., 2012). Although all three tissues (root, leaves and flowers) used in our study did not show such strong tissue-specific splicing signatures, some other plant tissues might. For example, the transcriptomes of sexual tissues are substantially different from those of vegetative tissues, and anthers harbor the most diverged specialized metabolomes (Yang and Wang, 2013; Li et al., 2016). Future studies that include transcriptome data of much more fine-scaled tissue samples will provide new insights on this aspect.

We found that the AS resulted in transcripts with PTC, which is likely coupled with nonsense-mediated decay (NMD) for degradation is more conserved than the AS that do not generate PTC-containing transcripts in plants (Figure 3A). Consistently, among six ultra-conserved AS events across different plant species including the spreading earth moss, five produced PTC+ transcripts, indicating that PTC+ AS might be more important than it was previously thought. Previous studies showed that all human serine/arginine-rich (SR) genes and some SR genes in plants produce AS resulted in PTC+ transcripts (Kalyna et al., 2006; Lareau et al., 2007; Palusa and Reddy, 2010). Furthermore, the junction regions that contain PTC+ AS in numerous splicing factors (SFs) are ultra-conserved between different kingdoms and the loss of the ancient PTC+ AS in paralogs through gene duplications were repeatedly replaced by newly created distinct unproductive splicing (Lareau and Brenner, 2015). In the same line, our results showed that many of these PTC+ AS are likely functionally important and are consistent with the hypothesis that the unproductive splicing coupled with NMD can be a functional process that controls the abundance of active proteins at a post-transcriptional level.

Among all five plant species, the distance between the 5’/3’ nearest alternative splice sites (SS) and the authentic SS is the main determinant that distinguishes AltD/AltA from constitutive splicing (Figure 4 and Supplemental file 1). For a given spliced junction, the likelihood of AS decreases with increased distance between the authentic and nearest alternative SS (Figure 4A and B). Interestingly, similar patterns were also found in mammals, in which, the closer the alternative SS was to the authentic SS, the more likely it was used for AS (Dou et al., 2006; Rosenberg et al., 2015). Interestingly, the frequency of AltA also decreases with the increased distance between the authentic SS and nearest inter-AG dinucleotide. This result is consistent with the pattern found in humans in that only closely located AGs (< 6 nt) can efficiently compete with the authentic SS and the distance between branch site (BS) and the first downstream AG can affect the 3’ SS selection (Chiara et al., 1997; Chua and Reed, 2001). Although, the BS in plants is not well studied and BS was not identified in ~30% of junctions (Reddy, 2007), the similar effects of inter-AG distance on AltA in both plants and mammals indicates that the mechanisms of generating AS, at least for AltA, might be similar between these two kingdoms.

While the deep learning model for AltA achieved high precision and specificity among five species (AUC > 0.9), the models for AltD performed less well than those for AltA, although still performing much better than by chance (AUC < 0.8, Figure 4C and D, S13). This indicates that additional determinants that contribute to the regulations of AltD were not detected by our method. It is known that the mechanisms involved in AltD are more complex than AltA. For example, in both human and mouse, while both the presence and quantity of exon splicing enhancer (ESE) and exon splicing silencer (ESS) are important for generating AltD (Koren et al., 2007), AltA is mainly affected by the competition of closely located AG dinucleotide by a scanning mechanism for the downstream sequence of the BS-polypyrimidine tract (PPT) (Smith et al., 1989; Smith et al., 1993; Chiara et al., 1997; Chua and Reed, 2001). These results suggest that splicing regulatory elements (SREs) may play more important roles in the proper selection of alternative SS in AltD than AltA. This may explain why the junction size contributed more in the AltD model than in the AltA model (Supplemental file 1), since larger junction size might increase the likelihood of introducing intronic SREs. In our attempts to identify SREs, although a few candidate sequence motifs were identified using the enrichment analysis, none of them significantly contributed to the model predictions. Two non-exclusive possibilities may explain this failure. First, the identified motifs are not involved in splicing regulation processes, although their density was significantly different between constitutively and alternatively spliced junctions. Second, they might be involved in tissue-specific regulations of AS, which likely did not contribute to the overall AltD prediction based on all three tissues. Future studies using different approaches to investigate the alteration of AS by introducing millions of random hexamers into specific regions of a gene junction in a plant then measuring the consequences of splicing, may allow us to more reliably detect splicing regulators of AltD in plants.

Although we found both junction size and SS for IR junctions are different from the constitutively spliced junctions (Figure S10), the identified features did not improve the AS prediction from that occuring by chance, indicating some other key determinants for IR remain missing in the model. As the expression level of IR is usually low and therefore requires high sequencing depth for their detection (Figure S16), it is likely that the sequencing depth of the transcriptome data used in this study was not sufficient to detect all of the IR junctions. Therefore, many true IR junctions may not have been considered as IRs in our analysis, which reduced prediction precision and power.

For both AltA and AltD, their rapid evolution between closely related species were mainly due to variations in the key sequence determinants near the SS (Figure 5, S14 and S15) and the key sequence determinants such as distance to authetic SS and *cis*-elements (BS, PPT, UA-rich tract for AltA) are all located within intronic regions. Intron sequences diverge rapidly (Mattick, 1994; Hare and Palumbi, 2003), therefore, the process of which likely have contributed to the rapid gains and losses of AS among different lineages to produce species-specific AS profiles in plants. For example, a decreased distance between alternative SS and authentic SS as a result of a short deletion of the intron sequence could lead to a gain of AS at the junction, and as consequence is likely to be shared among different tissues. Consistently, in vertebrates, the mutations that affect intronic splicing regulatory elements (SREs) were shown to be the main factor that resulted in the dominant species-specific splicing pattern (Merkin et al., 2012). However, our data can not exclude the possibility that the species-specific trans-factors, such as SR protein family, which have distinct numbers of homologues among species (Figure S17) (Iida and Go, 2006; Isshiki et al., 2006; Ling et al., 2015), may have also contributed to the divergence of AS among different species (Ast, 2004; Barbosa-Morais et al., 2012).

We also investigated other factors that were hypothesized to affect AS evolution, such as gene duplication (GD), DNA methylation and transposable element (TE) insertion (Sorek et al., 2002; Su et al., 2006; Flores et al., 2012). However, with the exception of TE insertions, the effects of which were found to be species-specific, most of the tested factors did not show significant effects on the levels of AS conservation between closely related species (Figure S15 and Supplemental file 2). The species-specific effects of TE on the AS conservation were likely due to the different abundance of TE insertions in the genomes of different species (Hu et al., 2011; Tomato Genome, 2012; Slotte et al., 2013; Sierro et al., 2014), suggesting genomic composition of each species might also affect the evolutionary alteration of AS.

## CONCLUSIONS

We found that the AS profiles diverged rapidly in plants, which is largely due to rapid gains and losses of AS in each lineage, while a group of AS that generate PTC-containing transcripts is highly conserved among very distantly related plants. The alteration of a few key sequence determinants of AltA and AltD, all located in the intron region, contributed to the rapid divergence of AS among closely related plant species. These results provide mechanistic insights into the evolution of AS in plants and highlight the role of post-transcriptional regulation of a plant’s responses to environment interactions.

## MATERIALS AND METHODS

### Read mapping, transcripts assembly and abundance estimation

The raw sequence reads were trimmed using AdapterRemoval (v1.1) (Lindgreen, 2012) with parameters “––collapse––trimns––trimqualities 2−−minlength 36”. The trimmed reads from each species were then aligned to the respective reference genome using Tophat2 (v2.0.6) (Trapnell et al., 2009), with maximum and minimum intron size set to 50,000 and 41 bp, respectively. The numbers of uniquely mapped reads and splice junctions (SJs) mapped reads were then counted using SAMtools (v0.1.19) (Li et al., 2009) by searching “50” in the MAPQ string and “*N*” flag in the CIGAR string of the resulting BAM files. The uniquely mapped reads from each sample were sub-sampled into same sequencing depth (16 million) using SAMtools (v0.1.19) (Li et al., 2009). The mapping information and IDs of all download datasets deposited in Sequence Read Archive (SRA) (http://www.ncbi.nlm.nih.gov/sra) were listed in Table S1 and S2.

The transcripts of each species were assembled using Cufflinks (v2.2.0) (Trapnell et al., 2012) with the genome annotation as the reference. The open reading frame (ORF) of each transcript was analyzed using TransDecoder from TRINITY (v2.1.0) (Grabherr et al., 2011). To estimate the expression level of genes/transcripts, all trimmed reads were re-mapped to the assembled transcripts using RSEM (v1.2.8) (Grabherr et al., 2011). Transcripts per million (TPM) was calculated for each gene/transcript (Wagner et al., 2012). Only genes with TPM greater than five in at least one sample were considered as an expressed gene.

### AS detection

All AS analysis were based on splicing junctions obtained from the BAM files produced by Tophat2. To remove the false positive junctions that were likely due to non-specific or erroneous alignments, all original junctions were filtered based on overhang sizes greater than 13 bp, as suggested in (Ling et al., 2015). All filtered junctions were then used for AS identification and annotation using JUNCBASE v0.6 (Graveley et al., 2011). Due to the relatively low sequence depth of each individual sample of Brassicaceae RNA-seq data (Seymour et al., 2014) (Table S2), we merged the BAM files of each three replicates together and random subsampled 17 million (the lowest depth among all merged samples) unique mapped reads from each merged to avoid the heterogeneity of sequencing depth.

The percent spliced index (PSI) of each AS event, which represents the relative ratio of two different isoforms generated by the AS was calculated in each sample. PSI = (number of reads to inclusion isoform) / (number of reads to inclusion isoform + number of reads to exclusion isoform) as suggested in (Graveley et al., 2011). To avoid false-positives, PSI was calculated only for AS events that had a total read count equal or greater than ten.

### Identification of conserved exon-exon junctions (EEJs) and AS

We separately extracted 100 bp sequence from the flanking upstream exon and downstream exon of each junction that have mapped read to support, and combine each side of exon sequence (in total 200 bp sequence) to represent the EEJ. The sequences of all EEJs from two species were searched against with each other using TBLASTX (v.2.2.25) (Altschul et al., 1990) to find homologous relationships (Figure S18). A python script was used to filter the TBLASTX results based on the following requirements: (1) The gene pair contain the EEJs must be the orthologous gene pair between the two species; (2) the EEJ sequences between two species must be the best reciprocal blast hit based on the bit score; (3) at least 3 amino acid (aa) from both the flanking upstream exon and downstream exon sequence were aligned and (4) alignment coverage >= 60 bp, (5) E-value <1E-3.

We only consider an AS event to be conserved if the same type of AS was found on the conserved EEJs between two plant species.

### Identification of AS that generates premature termination codons (PTC)

The junctions related to each AS event were mapped back to assembled transcripts; only AS which related to junctions that mapped to two unique transcripts (had no structural difference except the AS region) were retained to avoid the situation where the sequence differences of the two transcripts resulted from multiple AS events. The transcript was considered to have a PTC if the stop codon of the longest ORF is at least 50 nucleotides upstream of an exon-exon boundary (the 50 nucleotides rule) (Nagy and Maquat, 1998). The PTC-generating AS events are defined as only one of the resulting transcripts contain PTC.

### One-to-one orthologous gene identifications and gene family size estimation

One-to-one orthologous gene pairs were predicted based on pair-wise sequence similarities between species of the corresponding dataset. First, we calculated the sequence similarities between all protein-coding genes using BLASTP for the selected species and filtered the results based on E-value less than 1E-6. Second, we selected the groups of genes that represent the best reciprocal hits that are shared among all species from the corresponding dataset.

For calculating the gene family size, we first defined gene families among different species by using a similarity-based approach. To do so, we used all genes that were predicted from the respective genomes of each species. In brief, all-vs-all BLASTP was used to compare the sequence similarity of all protein coding genes, and the results were filtered based on the following criteria: E-value less than 1E-20; match length greater than 60 amino acids; sequence coverage greater than 60% and identity greater than 50%. All BLASTP results that remained after filtering were clustered into gene families using the Markov cluster algorithm (mcl) (Enright et al., 2002). The gene family size for a species is represented by the number of genes of this species within the corresponding gene family.

### Correlation and clustering

For the pairwise comparison of AS, Spearman correlation and binary distance was applied to the PSI data (0.05 < PSI <0.95 in at least one sample) and binary data (all genes that had no AS in all four species were excluded), respectively. A non-parametric correlation was selected for PSI level because of its bimodal nature distribution (0 and 100). For the pairwise comparison of GE, Pearson correlation was applied to log_2_ (TPM+1) of expressed genes to avoid infinite values.

The R package “pvcluster” was used for clustering of samples with 1,000 bootstrap replications. When we clustered and performed principal-component analysis (PCA) of gene expression, the TPM values were normalized by GC% (EDASeq package in R) and TMM (the trimmed mean of M-values).

### Identification of possible alternative splice sites (SS) and regulatory sequences

The 5’ and 3’ splice site including 5 bp up and downstream sequences of all EEJs were used as the positive dataset, while the sequences extracted using the same method for all inter-GT (for 5’ splice site) and inter-AG (for 3’ splice site) within junction regions were used as background dataset. The putative SS motifs (6-mer) of both 5’ and 3’ SS were separately identified using Homer V3.12 (Heinz et al., 2010) and only motifs present in at least 5% of total positive sequences and *P*-value<1E-20 were kept. The appearance of putative SS was identified using scanMotifGenomeWide, a Perl script included in the Homer toolkits and only sequence regions with match score >2 were kept.

Homer was also used to identify the putative regulatory intronic and exonic sequence motifs (6-mer) of AltD, AltA and IR. The 50 bp up and downstream sequence of 5’ SS was regarded as exonic and intronic sequence and vice versa for 3’ SS. For AltD and AltA, the related sequences of EEJs with AS were used as the positive dataset, while 10,000 related sequences of EEJs without AS by random selection (due to a large number of sequences) were used as background dataset. The enriched motifs in the positive dataset were regarded as splicing enhancers, while the enriched motifs in the negative dataset were considered as splicing silencers. For IR, the related sequences from both splice sites of EEJs with IR were used as the positive dataset and the same sequences from EEJs without IR were used as background dataset. The conserved motifs between species were identified using compareMotifs, a Perl script included in the Homer toolkits and only one mismatch was allowed. To identify PPT, UA-rich tracts and branch site (BS) of each EEJ, we used the Perl scripts from Szczesnia et al. (Schwartz et al., 2008; Szczesniak et al., 2013) To estimate the effect of each putative sequence motif, PPT and UA-tracts, we calculated the AS frequency of EEJs containing or not containing the motif/tract (Rosenberg et al., 2015). Then for each motif/tract, the log_2_ odds ratio (effect size) with and without the motif/tract were calculated to quantify to what extent the presence of the motif/tract increases or decreases the AS frequency compare to its absence: 

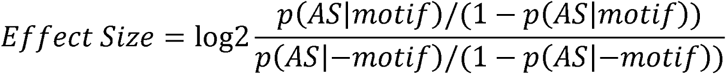

### Deciphering the splicing codes and AS conservation using deep learning algorithm

To investigate which sequence determinants contributed to the AS in plants, we constructed multi-layer feed-forward artificial neural networks using H_2_O’s deep learning algorithm (‘h2o’ package) in R 3.0.2 (R Development Core Team 2013). For each AS type, a matrix was created based on the information of all EEJs that contain the AS (only that single event) and other EEJs within the same gene. The AS status (either AS or constitutive) was considered as output and the features that were known to be associated with splicing recognition and regulation in eukaryotes (Lewandowska et al., 2004; Kandul and Noor, 2009; Rosenberg et al., 2015) (listed in Supplemental file 1) were used as input for training the model. To reduce the background noise, we removed the EEJs which were supported by less than five reads on average. In addition, because the number of constitutively spliced EEJs in all cases is much larger than alternatively spliced EEJs, we randomly selected the same number of constitutive spliced EEJs as alternative spliced EEJs and combined them together with all alternative spliced EEJs as the full dataset (50% precision by chance). To train and test the deep neural networks (DNN), the full dataset was randomly split, which 60% of data were used for training, 20% used for validation and the other 20% was reserved for testing. We trained for a fixed number (10,000) of epochs or stopped the training once the top 10 model were within 1% of improvement, and selected the hyper-parameters that gave the optimal AUC performance on the validation data. The model was then retrained using these selected hyper-parameters with the full dataset.

Using the similar approach, we constructed the model for AS conservation. For each AS type, a matrix was created based on the information of all orthologous EEJ pairs between two species that contain the AS in at least one species. To reduce the background noise, any EEJ with multiple AS types, low number of support reads (less than five) or orthologous EEJ pair have different AS types were removed. The conservation levels (conserved, lost or gained in the other species) were used as the output of the model and the difference of features that were known to be important to AS and AS conservation (Su et al., 2006; Kelley et al., 2014; Li et al., 2014; Lambert et al., 2015; Rosenberg et al., 2015) (listed in Supplemental file 2) between two species were used as input to train the model. Yass v1.15 (Noe and Kucherov, 2005) was used to align the SS’ flanking sequences (combined 50 bp upstream and downstream sequences of 5’/3’ splice site, 100 bp in total) of each orthologous EEJ pair, the similarity was calculated as: (length of alignment - number of gaps - number of mismatches) / (total sequence length). To reduce the bias from different transition types in the dataset (much higher proportion of loss/gain than conserved AS), the data used to train the model was selected as the ratio of 1:1:1 for conserved, lost and raised situations (33.3% precision by chance). Due to the rather small sample size of conserved AS, the model based on the same original data may differ as the randomly selected data of AS lost/raised were different each time. Therefore, the model construction process was repeated 10 times and the models that achieved the highest AUC for the complete dataset were considered.

## Accession Numbers

The ID of all RNA-seq data that are deposited or downloaded from NCBI short reads archive (SRA) database for generating the results in this study is listed in Table S1.

## ACKNOWLEDGMENTS

We thank Danell Seymour and Daniel Koenig for providing the methylation data, Michal Szczesniak for providing the Perl scripts for finding UA tracts.

### SUPPLEMENTAL MATERIAL

**Supplemental Figure S1.** The distribution of different types of alternative splicing (AS) events in (A) *A. thaliana*, (B) *G. max*, (C) *S. lycopersicum*, (D) *N. attenuate*.

**Supplemental Figure S2.** Species-specific clustering of alternative splicing (AS) among different plant species.

**Supplemental Figure S3.** Conservation of gene expression (GE) in eudicots.

**Supplemental Figure S4.** Comparative profiling of conserved junctions and alternative splicing in eudicots.

**Supplemental Figure S5.** Comparative profiling of conserved junction and alternative splicing (within one-to-one orthologues) in three Brassicaceae species.

**Supplemental Figure S6.** The transition spectrum among different types of AS between species pairs.

**Supplemental Figure S7.** The proportion of AS that generates PTC (PTC+) or not (PTC-) in *N. attenuata*, tomato, *A. thaliana* and soybean.

**Supplemental Figure S8.** The probability of DNA bases surrounding SS with different AS types compared to regular SS in five plant species.

**Supplemental Figure S9.** The complete linkage hierarchical clustering of SS motifs among different plant species.

**Supplemental Figure S10.** The determinants of intron retention (IR) in plants.

**Supplemental Figure S11.** The effect of UA-rich, polypyrimidine tract (PPT) and branch site (BS) on alternative acceptor (AltA) and intron retention (IR) in plants.

**Supplemental Figure S12.** The effect size of conserved 6-mer motifs and features identified in (A) alternative 5’ donor site (AltD) and (B) alternative 3’ acceptor site (AltA) between species pairs in Solanaceae and Brassicaceae.

**Supplemental Figure S13.** The area under the curve (AUC) of deep learning models using different key features of (A) alternative 5’ donor (AltD) and (B) alternative 3’ acceptor (AltA) in tomato, *A. lyrata* and *C. rubella*.

**Supplemental Figure S14.** Differences in the *cis*-regulatory elements affect the turnover rate of (A) AltD and (B) AltA between plant species.

**Supplemental Figure S15.** Factors that affect the rapid turnover of AS between plant species.

**Supplemental Figure S16.** The composition of AS types detected in leaf and root samples of *N. attenuata* with different sequencing depth.

**Supplemental Figure S17.** Phylogenetic tree of SR and SR-like genes in moss and four eudicots.

**Supplemental Figure S18.** The diagram showing the process of identifying conserved exon-exon junctions (EEJs) between two species.

**Supplemental Table S1.** RNA-seq coverage and alignment statistics of the four eudicots

**Supplemental Table S2.** RNA-seq coverage and alignment statistics of Brassicaceae

**Supplemental file 1:** This file contains two excel sheets that show the relative importance of each factor that contributes to the splicing model of AltD and AltA in five different species. SS: splicing site; BS: branch site; PPT: polypyrimidine tract. The top five contributors to each model were highlighted in red.

**Supplemental file 2:** This file contains two excel sheets that show the relative importance of each factor that contributes to the AS conservation of AltD and AltA among closely related species. SS: splicing site; BS: branch site; PPT: polypyrimidine tract. The top five contributors to each model were highlighted in red.

